# An oscillopathic approach to developmental dyslexia: from genes to speech processing

**DOI:** 10.1101/108704

**Authors:** Miguel Jiménez-Bravo, Victoria Marrero, Antonio Benítez-Burraco

**Affiliations:** Department of Philology, UNED, Madrid, Spain; Department of Philology, University of Huelva, Huelva, Spain

**Keywords:** Developmental dyslexia, brain rhythms, candidate genes, speech sounds, reading, spelling

## Abstract

Developmental dyslexia is a heterogeneous condition entailing problems with reading and spelling. Several genes have been linked or associated to the disease, many of which contribute to the development and function of brain areas that are important for auditory and phonological processing. Nonetheless, a clear link between genes, the brain, and the symptoms of dyslexia is still pending. The goal of this paper is contributing to bridge this gap. With this aim, we have focused on how the dyslexic brain fails to process speech sounds and reading cues. We have adopted an oscillatory perspective, according to which dyslexia results from a deficient integration of different brain rhythms during reading/spellings tasks. Moreover, we show that some candidates for this condition are related to brain rhythms. This approach should help gain a better understanding of the aetiology and the clinical presentation of developmental dyslexia, but also achieve an earlier and more accurate diagnosis of the disease.

## 1. Introduction

Developmental dyslexia is a neurobiological condition entailing reading and spelling problems, in spite of normal intelligence and adequate environmental feedback from parents, teachers, and caregivers (Démonet et al., 2004). This is the most common learning disability in children, with a prevalence of 5 to 17.5% among school-aged children (Shaywitz, 1998). Dyslexia is more frequent in boys, with a sex-ratio between 1.35:1 and 2.76:1 (Rutter et al., 2004; Quinn and Wagner, 2015), and with higher ratios usually found in clinical-referred samples (Finucci and Childs, 1981) and among the most severe cases (Hawke et al., 2009). Dyslexia is less prevalent in languages with shallow orthographies, like Spanish, which usually involve univocal links between graphemes and phonemes (Wagner et al., 2005); (Davies et al., 2007). Nonetheless, this condition has been found in all languages and in all writing systems (Paulesu et al., 2001).

Developmental dyslexia has a genetic basis and results from broader cognitive deficits mostly impacting on reading acquisition and processing (Darki et al., 2012; Skeide et al., 2015). Accordingly, compared to controls, dyslexics show poorer digit span skills, reduced abilities for word formation, and slower automatic naming abilities (Wagner and Torgesen, 1987). Also, core perceptual capacities seem to be impaired in them. Hence, dyslexic children fail to process quick changes in sound features that are important for speech understanding, like formant transitions or amplitude envelope modulations (Tallal, 1980). Finally, they exhibit poorer motor control and impaired balance (Nicolson and Fawcett, 1990; for a review see Rochelle and Talcott, 2006). This heterogeneous clinical profile has resulted in different hypothesis about the etiopathogenesis of the disorder, and ultimately, in different classifications of the disease. The phonological deficit theory is perhaps the most renowned account of the causes of dyslexia (e.g. Vellutino, 1979; Tallal, 1980; Stanovich, 1988; Liberman et al., 1989; Snowling, 2000; Ramus, 2001; Vellutino et al., 2004). Under this view, dyslexia results from a deficit in phonological processing (that is, the ability to access phoneme representation from auditory cues), which in turn results from the dysfunction of the perisylvian cortex (Galaburda et al., 1985). Nonetheless, according to others dyslexia may be caused by a broader deficit in the ability to process rapidly changing stimuli, either auditory or visual, or both (Tallal, 1980; Eden et al., 1994). Other researchers have claimed that dyslexia may result from a failed synchronization and integration of visual and auditory input, hence the magnocellular account of this condition (the magnocellular pathways of the thalamic medial and lateral geniculate nucleus function as relay station for rapidly modulated input) (Galaburda et al., 1985; Breznitz and Meyler, 2003; Sela, 2012). Finally, the cerebellar deficit theory of dyslexia supports the view that this condition is caused by a deficit in non-verbal sensory-motor integration, in which the cerebellum plays a key role (Nicolson et al., 2001).

Usually, an L-type dyslexia and a P-type variant of the disease are posited. L-type (or phonological, or dysphonetic) dyslexia entails problems for sounding out words through grapheme-phoneme mapping, whereas P-type (or surface, or dyseidetic) dyslexia mostly involves problems for recognizing whole-word visual configurations (Bakker, 1973; Boder, 1973; Castles and Coltheart, 1993; Coltheart et al., 1993). The clinical diagnosis and the therapeutic management of dyslexia is hindered by its frequent co-morbidity with other conditions, including mathematic disorder (Dirks et al., 2008; Willcutt et al., 2013), attention-deficit/hyperactivity disorder (ADHD) (Willcutt et al., 2005; Sexton et al., 2012), speech-sound disorder (SSD) (Pennington and Bishop, 2009), specific language impairment (SLI) (McArthur et al., 2000; Bishop and Snowling, 2004), and developmental coordination disorder (Kaplan et al., 1998). It seems that if we want to achieve a clear categorization of dyslexia and a more accurate view of the origins of the disease, we need a better account of how the involved biological factors contribute to the disease.

We have structured our paper as follows. First, we provide a general overview of language deficits in dyslexia. Next, we focus on the dyslexic brain and give a brief account of the brain areas that are structurally and/or functionally impaired in dyslexics. Third, we focus on the abnormal brain oscillatory activity observed in dyslexic patients, and discuss putative links between this oscillopathic profile and the speech/reading deficits observed in them. Then, we move to the genes and provide a brief account of the candidates for this condition, with a focus on those that might help explain the observed oscillopathic profile of dyslexics. We will conclude by claiming that this approach should help clarify the neurocognitive profile of the affected people, but also achieve better therapeutic strategies.

## 2. The neurocognitive profile of dyslexics

Dyslexics usually achieve low performance scores in tasks evaluating three different, but related, aspects of phonological processing: phonological awareness (that is, the ability to perceive, evaluate, and manipulate the sounds of speech), automatized naming (that is, the ability to retrieve and name items from a list), and verbal short-term and working memory (which enables to storage, manipulate, and repeat items from a list) (Wagner and Torgesen, 1987). The precise nature of the phonological deficit underlying dyslexia is not clear (for discussion see Ramus and Szenkovits, 2008; Szenkovits et al., 2016). It might result from degraded or weak phonological representations (Bradley and Bryant, 1983; Elbro et al., 1998; Griffiths and Snowling, 2002; Noordenbos and Serniclaes, 2015), from poor categorical perception (Manis et al., 1997; Adlar and Hazan, 1998), or even from poor auditory processing, particularly of rapidly changing short auditory stimuli, as occurs in formant transitions (Tallal, 1980, Snowling, 2000). Some authors have claimed instead that phonological representations are in fact over-represented in dyslexics, because they encode allophonic (i.e. non-phonological) features too (Serniclaes et al., 2004; Bogliotti et al., 2008; for a review see Noordenbos, 2013). Phonological problems in dyslexia have been hypothesised to result as well from inaccurate storage and retrieval of otherwise intact phonological representations (MacSweeney et al., 2009; Kovelman et al., 2012; Ramus and Ahissar, 2012; Boets et al., 2013). Finally, some authors have claimed that dyslexics might alter subsequent phonological processing because of a deficit in the left auditory cortex (Molinaro et al., 2016). In truth, dyslexics score like controls in all phonological tasks except when short-term memory load increases (de Jong, 1998; Isaki et al., 2008; Laasonen et al., 2012).

As noted above, not only auditory but also visual impairment has been observed in people with dyslexia (Dunlop, 1972; Lovegrove et al., 1980). Besides de ability to correlate graphemes and phonemes, reading also entails the ability to capture visual elements at a glance –typically letters– and to parse them in strings (Pelli and Tillman, 2007). Several studies have concluded that dyslexics suffer (as well) from a deficit in sequential/serial mechanisms involved in visual searching (Casco and Prunetti, 1996), and ultimately, from a visual-attentional deficit involving the magnocellular system (Vidyasagar and Pammer, 1999). Likewise, motor behaviour is also altered in people with dyslexia, impacting on equilibrium (Nicolson and Fawcett, 1990; Wimmer et al., 1999; Rochelle and Talcott, 2006), ocular movement (Eden et al., 1994; Biscaldi et al., 2000; Ram-Tsur et al., 2006), coordination, and speed motor tasks (Fawcett and Nicolson, 1999; Stoodley et al., 2006).

The disparate symptoms of dyslexia, involving the perceptual, cognitive, and motor domains, demands an integrative account of the etiopathogenesis of this condition. The cerebellar dysfunction hypothesis highlights the role of the cerebellum in the automatization of the varied processes underlying reading (Nicolson and Fawcett, 1990; Fawcett et al., 1996; Nicolson et al., 2001; for a review see Stoodley and Stein, 2011). Likewise, “sensory theories” of dyslexia argue for a wider interpretation of the phonological deficit theory, relying on auditory, visual, and motor deficits (Goswami, 2014). Nonetheless, as noted by Nimmrich, Draguhn, and Axmacher (2015, p. 272), “the increasing knowledge about patterned network activity suggests that dysfunctions of the nervous system cannot be understood without analysing the respective (oscillating) network activity patterns”. Accordingly, the analysis of brain rhythmicity during reading and spelling tasks by dyslexics should help clarify the etiopathogenesis of this condition. Before delving into the oscillopathic nature of dyslexia, we will briefly review the evidence pointing to structural and functional anomalies in the brain of the affected people.

## 3. The dyslexic brain

The complex neurocognitive profile of dyslexia boils down to the impairment of the different neural subsystems involved in reading, in particular, the auditory-phonological, the visuo-magnocellular, and the motor/cerebellar subsystems (Danelli et al., 2013). Structural disturbances found in the dyslexic brain include ectopias and focal microgyria in perisylvian regions (Galaburda and Kemper, 1979; Galaburda et al., 1985), abnormal layer organization in the lateral and medial geniculate nucleus of the thalamus (Livingstone et al., 1991; Galaburda et al., 1994), defects in the cerebellum (Finch et al., 2002; Rae et al., 2002), and anomalies in the corpus callosum (Rumsey et al., 1996; Robichon and Habib, 1998). Also, the asymmetry of the planum temporale and of Heschl's gyri observed in controls has not been reported in dyslexics (Galaburda et al., 1985; Humphreys et al., 1990; Altarelli et al., 2014). In turn, an abnormal pattern of neuronal size asymmetry has been found in them in the primary visual cortex (Jenner et al., 1999). Likewise, a reduction of grey matter in left frontal, temporoparietal, occipital areas has been observed in patients with dyslexia, as well as changes in grey matter volumes in other brain areas that correlate with specific cognitive deficits, like reduced phonological awareness, slow automatized naming, impaired magnocellular-dorsal processing, and deficits in auditory attention shifting (Jednoróg et al., 2014; Xia et al., 2016). The reduction of grey matter volumes seems to affect women more than men, particularly regarding the sensory and motor cortices (Evans et al., 2014). Dyslexia also entails changes in white matter volumes and white matter integrity in the left hemisphere, particularly, in the arcuate fasciculus, which correlates with the phonological and reading deficits exhibited by dyslexics (Vandermosten et al., 2012; Saygin et al., 2013).

Regarding functional anomalies, PET and fMRI studies have revealed an underactivation of the posterior areas of the temporoparietal and occipitotemporal regions of the left hemisphere during reading tasks by dyslexics, which is suggestive of problems for accessing phonological representations, as well as a compensatory overactivation of the same regions in the right hemisphere (Rumsey et al., 1997; Brunswick et al., 1999; Paulesu et al., 2001, 2014; Pugh et al., 2001; Shaywitz et al., 2002). According to Paulesu et al. (2014), this region in the left hemisphere comprisses three sub-areas that are important for reading: dorsally, a lateral inferotemporal multimodal area, associated with the integration of orthography and phonology; ventrally, the so-called “visual word form area” (VWFA), involved in quick visual processing of familiar strings of letters and in whole-word recognition; and finally, an intermediate area, located in the inferior temporal gyrus.

Importantly, differences in connectivity patterns during reading tasks have been also observed in dyslexics. Specifically, in children with dyslexia the strong connectivity found in controls between the left occipitotemporal area (processing visual information) and the left inferior frontal cortex (processing articulatory and phonological information) is not observed (van der Mark et al., 2011; Olulade et al., 2015). More generally, whole-brain functional connectivity analyses have revealed generalized changes in connectivity patterns in the brain of dyslexics, which affect the normal interconnection of the areas comprising the default-mode network involved in reading, but also its connection with other subsidiary networks, like the executive network (Finn et al., 2014; Schurz et al., 2015; Zhao et al., 2016). Of particular interest are the reported anomalies found in dyslexics in the magnocellular pathway, which provides an input to the dorsal stream encoding movement and rapid changes in the visual field (Lovegrove et al., 1986; Stein and Walsh, 1997). These anomalies seemingly impact on visual processing, much in line with the deficits observed in auditory domain (Farmer and Klein, 1995; Stein and Walsh, 1997; Hari and Renvall, 2001; McLean et al., 2011), accounting for the visual-attentional deficit found in people with dyslexia (Steinman et al., 1997; Hari et al., 1999; Vidyasagar and Pammer, 1999; Facoetti et al., 2003; for a review see Wright et al., 2012).

Two additional lines of research are improving our understanding of the dyslexic brain. On the one hand, comparative functional imaging studies with users of different writing systems (logographic vs. alphabetic, shallow vs. deep alphabetic orthographies, etc.) are helping identify the core brain mechanisms and areas involved in reading, as well as those that are script-specific (Paulesu et al., 2000, 2001; Siok et al., 2008; Hu et al., 2010; Liu et al., 2013; Pollack et al., 2015; Qi et al., 2016). For instance, both shallow and deep orthographies result in the underactivation of regions within the left occipitotemporal cortex of dyslexic people.

Nonetheless, non-transparent orthographies give rise as well to the underactivation of the left frontal gyrus triangularis and the right superior temporal sulcus, and the overactivation of the left anterior insula; in turn, transparent orthographies result in the underactivation of the left frontal gyrus pars orbitalis and pars opercularis, and the overactivation of the left precentral gyrus (Martin et al., 2016). On the other hand, animal models of the disease are helping refine our view of the structural and functional problems associated with dyslexia. For instance, surgical induction of cortical microgyria and ectopias in rats have proven to impact on cortical connectivity, particularly, on cortico-thalamic circuits (Galaburda, 1999; Rosen et al., 2000), and recapitulate some of the symptoms of dyslexia, most notably, the impairment in auditory capacities (Fitch et al., 1994; Herman et al., 1997; Peiffer et al., 2004), and the learning deficits, including problems with working memory (Rosen et al., 1995; Waters et al., 1997; Balogh et al., 1998; Hyde et al., 2000). Likewise RNA interference (RNAi) of some candidate genes for the disease, like *DYX1C1* or *DCDC2*, impacts on neuronal migration and results in cortical malformations that resemble the structural anomalies found in the brain of dyslexics; additionally, it gives rise to a reduced processing capacity of complex sounds and spatial cues (Threlkeld et al., 2007; Burbridge et al., 2008). According to Ramus (2006), these evidences give support to the auditory and magnocellular hypothesis of dyslexia, and provide a neural basis to the occurrence in the affected people of a phonological deficit accompanied by additional sensorimotor problems.

## 4. Rhythms in the dyslexic brain

Neuronal oscillatory activity has been known for decades (Berger, 1929), but only recently the role of cortical rhythms in different cognitive and behavioural processes has been properly acknowledged and examined (for a review see Buzsáki, 2006). Regarding language, speech processing has been a focus of particular attention. Interestingly, as discussed below in detail, the anomalous patterns of neuronal oscillations found in the dyslexic brain suggest that this condition can be successfully construed as an oscillopathic disease, much in the line of language deficits in schizophrenia (SZ) or autism spectrum disorders (ASD) (Benítez-Burraco and Murphy, 2016; Murphy and Benítez-Burraco, 2016).

The auditory cortex performs a multi-time resolution analysis and integration of sounds. In a nutshell, neuronal oscillations in different frequency bands (delta, theta, alpha, beta and gamma) parse the sound input at the proper timescale (syllabic, phonemic, etc.), whereas top-down signals from the frontal areas modulate the phase of slow-rate brain oscillations in the auditory cortex and help detect the edges of speech envelope (Poeppel, 2003; Ghitza, 2011; Gross et al., 2013; Park et al., 2015). The most detailed account of speech processing in terms of brain rhythms is Poeppel’s Asymmetric Sampling in Time theory, which has crystallized in his Multi-Time Resolution Model (Poeppel, 2003; Poeppel et al., 2008). In brief, each hemisphere processes speech sounds at different time scales. The left hemisphere focuses on short temporal windows (approx. 20-80 ms), which are optimal for the analysis of phonemes and subsegmental elements, like distinctive features, and which correlate with gamma oscillations. In turn, the right hemisphere focuses on long integration windows (approx. 150–300 ms), which enable to extract syllabic cues and which correlate with delta and theta oscillations. Further cross-frequency coupling of oscillations at delta, theta, beta, and gamma bands, which follows a hierarchical pattern, helps achieving a successful speech parsing and recognition. As a consequence, the sound stream becomes segmented into discrete chunks that constitute the basic coding elements for subsequent neuronal computation, in particular, for accessing the semantic and syntactic levels (for details see Giraud and Poeppel, 2012, Ghitza et al., 2012, and Gross et al., 2013).

Altered patterns of brain oscillations have been reported in dyslexics for a long time (Flynn et al., 1992; Klimesch et al., 2001). Below we provide with a detailed account of the oscillatory anomalies found in the different bands of interest for speech/reading processing (see table 1 for a summary).

**Table 1.**
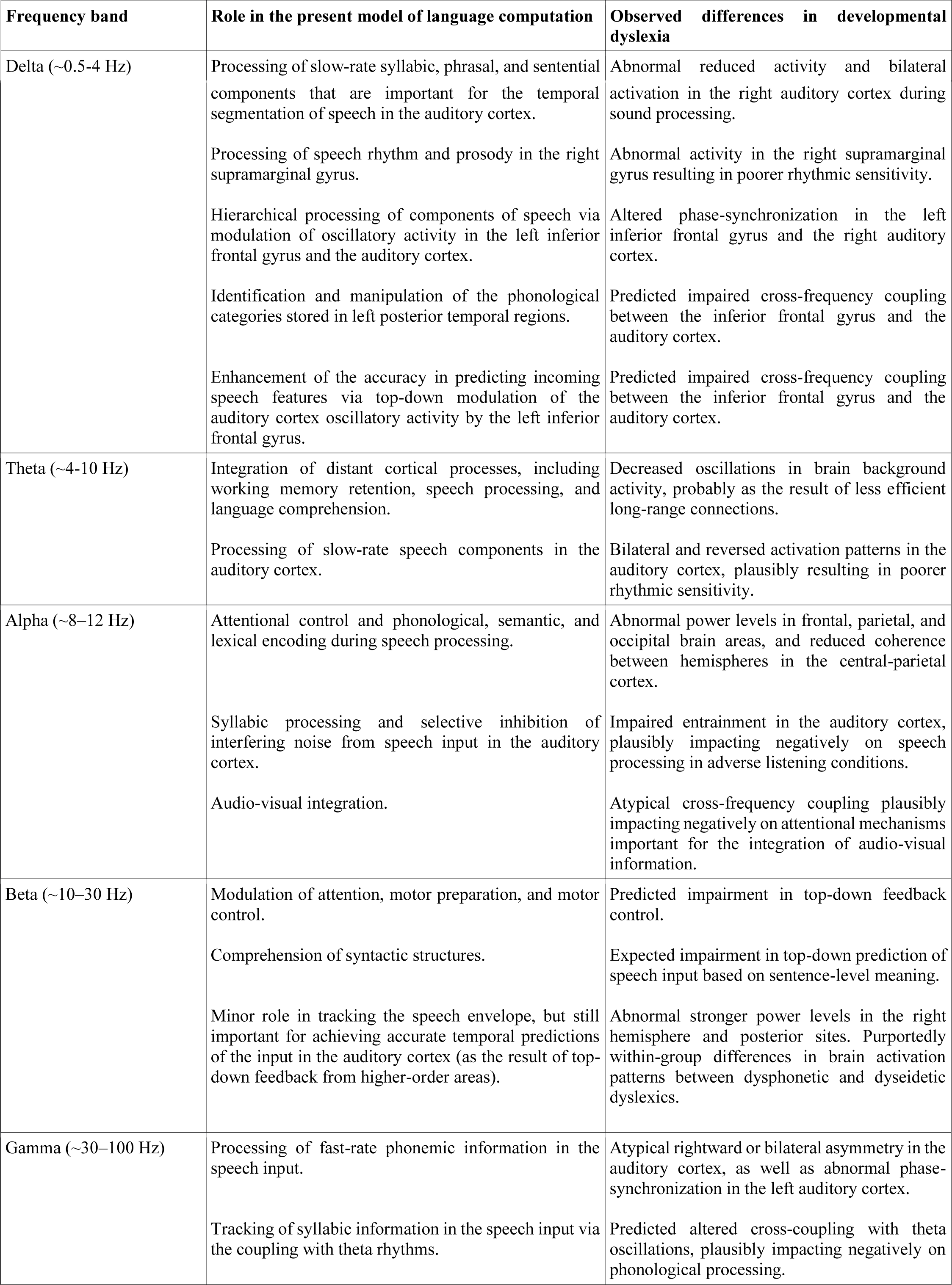
An outline of oscillopathic findings in dyslexia

### Delta (~0.5-4 Hz)

In the delta range, the dyslexic brain exhibits a more bilateral activity, as well as a decreased phase-synchronization when processing sounds, which contrasts with the rightward asymmetry and the strong phase-synchronization typically found in non-impaired readers. Using amplitude-modulated white noise, Hämäläinen and colleagues (2012) found less entrainment in the right auditory cortex of dyslexics at 2 Hz, but normal phase-synchronization of theta, alpha, and beta frequencies. In an EEG experiment, Soltész and colleagues (2013) observed weaker entrainment of the right auditory cortex of patients when they processed tone streams delivered at 2 Hz. Interestingly, they also found a reduced contingent negative variation, which is a component related to the anticipatory behaviour of the brain (see Arnal et al., 2015 for details). Consequently, Soltész and colleagues hypothesised that a link might exist between reading performance and neuronal anticipatory phase-synchronization in the delta range. Abnormal delta oscillations in the auditory cortex have also been found in dyslexics during the processing of speech sounds (Molinaro et al., 2016; Power et al., 2016); but see Lehongre et al., 2013; Lizarazu et al., 2015 for opposite results).

Abnormal delta response in dyslexia has been related to problems for correctly processing slow-rate speech information, in particular, rise-times in amplitude envelope, which correspond to syllable stress patterns (Goswami et al., 2013). Thus, the temporal segmentation of continuous speech via cortical phase-synchronization observed in non-impaired readers is atypically perfomed in dyslexics (Power et al., 2013, 2016; Doelling et al., 2014). Interestingly, differences between dyslexic and non-dyslexic readers in the 2 Hz band response have been found in the right supramarginal gyrus (Cutini et al., 2016). This brain region is involved in the processing of speech rhythm (Geiser et al., 2008), and speech prosody (Sammler et al., 2015), and is part of the dorsal stream for speech processing which is thought to be impaired in dyslexics. Nonetheless, because of the role of delta oscillations in hierarchical cross-frequency coupling, the atypical entrainment in the delta range observed in dyslexics might also alter the integration of speech information between low-level and high-level areas within the auditory cortex, but also between the auditory cortex and other cortical areas. Accordingly, Molinaro and colleagues (2016) found altered phase-synchronization in the delta range not only in the right auditory cortex of dyslexics, but also in the left inferior frontal gyrus (IFG). This brain area is involved in high-order computations as part of a larger language network (see Hickok and Poeppel, 2007; and Hickok, 2009 for details), and shows altered connectivity patterns in people with dyslexia (van der Mark et al., 2011). Specifically, this impaired entrainment of delta range oscillations in the right auditory cortex, and its concomitant effect on the oscillatory activity of the left IFG, might impact negatively on the correct identification and manipulation of the phonological categories stored in left posterior temporal regions. Phonological representations have been suggested to be intact in dyslexics, who would be instead unable to properly access them because of deficient top-down interaction between the left IFG and the right and left auditory cortices (see Boets et al. 2013 and Ramus, 2014 for discussion). This deficient top-down interaction has been recently identified as an abnormal modulation of delta and theta phase by the IFG, which results in a decreased phase-synchronization to continuous speech in the primary auditory cortex, ultimately impacting on the predictability of incoming speech (Park et al., 2015). Nonetheless, it seems now that an impaired bottom-up connection between perceptual (auditory cortex) and phonological (left IFG) processing regions should be expected too (Molinaro et al. 2016).

### Theta (~4-10 Hz)

Evidence of an abnormal phase-synchronization in the theta range in the auditory cortex of dyslexics is not conclusive (see Hämäläinen et al. 2012; Lehongre et al., 2013; Poelmans et al., 2012 for discussion). Increased bilateral, or reversed asymmetric patterns of activation for the theta range have been found in affected people compared to controls (Spironelli et al., 2008; Lizarazu et al. 2015). Likewise, decreased theta range oscillations have been observed in dyslexics in non-synchronized background brain activity (Fraga González et al., 2016; De Vos et al., 2017; although see Babiloni et al., 2012; Pagnotta et al., 2015 for opposite findings). This latter finding might be indicative of an altered interaction between the auditory cortex and the background activity of the cortex (De Vos et al., 2017) and/or of atypical brain connectivity patterns (Fraga González et al., 2016). The reason is the known role of slow oscillations such as theta in the integration of computations involving distant cortical regions (Buzsáki and Draguhn, 2004), which is important for working memory retention (von Stein and Sarnthein, 2000), speech processing (Luo and Poeppel, 2007), and language comprehension (Bastiaansen et al., 2008).

### Alpha (~8–12 Hz)

Anomalous cortical oscillations in the alpha band have been also reported in people with dyslexia. In a reading task involving words and pseudowords, Klimesch and colleagues (2001) found altered alpha power in frontal and central regions compared to controls, which points to attentional problems during the encoding of words. Likewise, lower amplitude of the alpha oscillations at parietal, occipital, and temporal sites seems to correlate with lower scores in pseudoword reading tasks (Babiloni et al., 2012). Additionally, dyslexics show reduced interhemispheric coherence in central–parietal areas when performing visuo-spatial attention tasks (Dhar et al., 2010). Overall, anomalies in the high–alpha band are expected to impact on phonological, semantic, and lexical processing, whereas anomalies in the low-alpha band are thought to impact on cortical arousal and vigilance (Babiloni et al., 2012). Interestingly, impaired phase-synchronization in the auditory cortex involving the alpha band has been recently reported in dyslexics, notably at 10 Hz (De Vos et al., 2017). This suggests that alpha oscillations may play some important role in syllabic processing too, together with slow-rate cortical oscillations. Also, they seem to be important for auditory selective inhibition, which enables to exclude noise interference from task-relevant information (Strauß et al., 2014). Accordingly, dyslexics perform worse than control in speech processing in adverse listening conditions (De Vos et al., 2017). Finally, atypical cross-frequency coupling involving the alpha band has been hypothesised to impact negatively on the attentional mechanisms responsible for integrating visual and auditory inputs, and ultimately, to account for some of the observed deficits in dyslexics (Klimesch, 2012). On the one hand, occipital alpha oscillations, one of the best-known rhythms arising from the visual system, are influenced by sound perception (Romei et al., 2012). On the other hand, larger alpha power over visual cortices is observed when attention is focused on the auditory component of an auditory-visual stimulus (Klimesch, 2012).

### Beta (~10–30 Hz)

Outside the auditory cortex, beta oscillations have been involved in the modulation of attention (Siegel et al., 2012), in motor preparation and control (Salmelin et al., 1995; Cheyne et al., 2012), and in syntactic comprehension (Bastiaansen et al., 2010; Lewis et al., 2016). Regarding the beta range, dyslexics exhibit an abnormal resting-state local efficiency (Dimitriadis et al., 2013), as well as stronger right versus left, and posterior versus anterior cortical asymmetries (Milne et al., 2003; Spironelli et al., 2008; Penolazzi et al., 2010). It has been claimed that stronger beta power in anterior brain regions is a hallmark of dysphonetic dyslexics, who experience problems with grapheme-to-phoneme conversions, whereas stronger beta signals in posterior regions of the brain are typical of dyseidetic children, who have problems for accessing the visual lexicon (Flynn et al. 1992, Milne et al., 2003). Interestingly, differences between young and adult subjects with dyslexia have been found regarding the abnormal beta oscillation pattern in the auditory cortex. Accordingly, an enhanced right hemisphere activity in the beta range has been found in teenagers and adults, but not in children (Lizarazu et al., 2015; De Vos et al., 2017; Power et al., 2016). Contrary to delta, theta, or gamma oscillations, beta rhythms seem to play a minor role in tracking the speech envelope, and consequently, in the etiopathogenesis of dyslexia (Power et al. 2016). Nonetheless, they may help achieve accurate temporal predictions of speech because of their coupling with delta oscillations in the auditory and motor cortices (Arnal et al., 2014; see also Park et al. 2015).

### Gamma (~30–100 Hz)

Atypical activation and synchronization of the gamma band have been attested in the auditory cortex of dyslexics compared to controls. Whereas non-impaired readers exhibit a leftward lateralization of the signal, dyslexics show a bilateral pattern of activation at 30 Hz (Lehongre et al., 2013; Lehongre et al., 2011), or even a rightward asymmetry at this frequency (Lehongre et al., 2011; Han et al., 2012; Poelmans et al., 2012; Lizarazu et al., 2015). This overactivation of the right auditory cortex has been hypothesised to reflect some compensatory mechanism aimed to deal with reading difficulties (see Paulesu et al., 2001; Shaywitz et al., 2002; Spironelli et al., 2006, 2008 for this view). Compared to controls, dyslexics also exhibit a stronger phase-synchronization in the left auditory cortex at high gamma frequencies (above 50 Hz), which might be indicative of the flooding of the auditory system with overdetailed spectrotemporal information, and ultimately, of a saturation of the theta-based auditory buffering capacity and of verbal working-memory (see Lehongre et al. 2011 for discussion). Some of these anomalies are not found in children with dyslexia. Accordingly, the stronger entrainment for the low (30 Hz) and high (60 Hz) gamma range is only observed in adults, whereas the rightward asymmetry for the gamma range observed in children becomes bilaterally distributed in adults (Lizarazu et al., 2015). These changes might reflect, at least in part, the prolonged effect of reading experience. Interestingly, in control subjects the phase-synchronization in the gamma range at 30 Hz correlates with cortical thickness asymmetries and patterns of cortical pruning in the auditory cortex; on the contrary, in dyslexics these structural and functional features correlates with phase-synchronization in the theta band at 4 Hz (Lizarazu et al., 2015). These results support the view that controls rely more on phonemes during reading tasks, whereas dyslexics depend more on syllabic units (see Lizarazu et al., 2015 for discussion).

## Overview

Giraud and Poeppel (2012) have argued for a crucial role of gamma oscillations in the etiopathogenesis of dyslexia. Under their view, dyslexics parse speech at frequencies that are slightly higher or slightly lower than the usual low gamma rate found in controls. In turn, this would result in phonemic units that are either undersampled (less acoustically detailed) or oversampled (too acoustically detailed, which is a burden for memory), in absence of major perceptual deficits. This phonological impairment would ultimately result in inaccurate grapheme-to-phoneme matchings.

Recent results by De Vos and colleagues (2017) show normal gamma band activity at 40 Hz in dyslexics. At the same time, as discussed above, the coupling of gamma and theta oscillations plays a key role in normal speech encoding and processing (see also Hyafil et al., 2015). Accordingly, gamma oscillations might play a basic role in cross-frequency coupling before the processing of the phonemic components of speech. Alternatively, as suggested by Goswami (2011), the main deficit in dyslexia might be an inefficient phase-synchronization of slow frequencies in the delta and theta ranges to prosody and syllable rates, respectively. According to her Temporal Sampling Framework hypothesis, this abnormal phase-locking would result in several other deficits that are typically found in dyslexics. Accordingly, their problems for correctly forming internal representations of aspects of speech rhythm would result in poorer rhythmic sensitivity, sluggish visual attention shifting, motor/cerebellar dysfunction, and magnocellular deficits, including altered auditory and visual integration. Also, the difficulties for noise exclusion might be explained in terms of a reduced perceptual sensitivity to amplitude and frequency modulation at lower rates (for details see Goswami, 2011).

Post-mortem examinations of the brains of affected people, but also animal models of the disease are helping refine this view. Accordingly, the ectopias and neuronal migration problems that are typically associated with dyslexia might disrupt specifically the normal organization of neurons within layers II and III of the cortex, which are responsible for low gamma and theta oscillations, and which temporally organize stimulus-driven spike trains coming from layer IV of the auditory cortex (Giraud and Ramus, 2013). This altered connectivity would prevent spike trains from layer IV from being properly transformed by gamma and theta oscillations. Therefore, the signal would be unable to reach the superficial layer in a read-out form that is appropriate for subsequent processing. As a consequence, by virtue of the hierarchical nesting of oscillations in the auditory cortex (Lakatos et al., 2005; Gross et al., 2013), an abnormal phase-synchronization to speech input at low frequencies may result in abnormal oscillations at higher frequencies, thus altering the proper encoding of speech at the phonemic level.

## 4. Candidate genes for dyslexia and brain rhythmicity

The number of genes related to developmental dyslexia has been growing over time (see Paracchini et al., 2016 for the most recent review). Interestingly, some of them are involved in brain rhythmicity and/or have been related to abnormal patterns of neuronal activity, like epileptiform behaviour. Although the available data are still scarce and fragmentary, we regard these genome-to-brain-to-language links as promising avenues of research to explain the processing deficits observed in this condition from an oscillopathic view (Figure 1). Below we provide a brief functional and biological characterization of the candidates for dyslexia with some known roles in brain rhythmicity.

Among the candidate genes resulting from association studies, as listed by Paracchini et al. (2016), one finds promising candidates for the oscillopathic profile of dyslexia. A late mismatch negativity (MMN) has been associated to rare variants in one intron of *DCDC2* and in the intergenic region between *DCDC2* and *KIAA0319* (locus DYX2) (Czamara et al., 2011). This MMN is a negative ERP component around 300-710 ms, which is indicative of differences between dyslexic children and age-matched controls when discriminating between complex auditory stimuli, like syllables and words, and which is originated in central-parietal areas of the right hemisphere (Hommet et al., 2009). *DCDC2* encodes a doublecortin domain-containing protein involved in neuronal migration in the cortex (Burbridge et al., 2008). *KIAA0319* encodes a membrane protein with several PKD repeats that plays a key role in the interaction between neurons and radial glial cells during neuronal migration (Paracchini et al., 2006; Velayos-Baeza et al., 2007). *CYP19A1* is also of interest. It encodes a cytochrome P450 protein with an aromatase activity that catalyses the formation of aromatic estrogens from androgens, which are hormones involved in neuronal plasticity that have been hypothesized to contribute as well to epilepsy (Fucic et al., 2009). Estrogen depletion by aromatase inhibition impacts on GABA synthesis and results in increased spine density and decreased threshold for seizures in the hippocampus (Zhou et al., 2007). CNVs of *CYP19A1* have been identified by array-CGH in patients with epilepsy (Kim et al., 2007). Finally, *ROBO1* has been found to regulate interaural interaction in auditory pathways (Lamminmäki et al., 2012). The gene is also a target of miR-218, which is significantly downregulated in the hippocampus of patients with mesial temporal lobe epilepsy (Kaalund et al., 2014). *ROBO1* encodes an axon guidance receptor that contributes as well to interneuron migration in the forebrain, and the development of ascending or descending axon tracts to or from the forebrain, specifically of thalamocortical axons, which modulate cognitive functions, consciousness and alertness (Andrews et al., 2006; Lopez-Bendito et al., 2007; Farmer et al., 2008; Dugan et al., 2011; Marcos-Mondejar et al., 2012). According to Wang et al. (2015), *ROBO1* was co-opted for vocalization in songbirds as part of the specialized motor song output nucleus, in which the gene is upregulated during critical periods for vocal learning. Interestingly for dyslexia, a splice variant of ROBO1, called ROBO1a, is highly enriched in the temporal auditory neocortex (Johnson et al., 2009).

Among the candidates for dyslexia resulting from GWAs, the most promising gene is *COL4A2*. It encodes the alpha 2 chain of type IV collagen, which is the major structural component of glomerular basement membranes; nonetheless, its C-terminal portion has also inhibitory and apoptotic activities, because it arrests the proliferation and migration of endothelial cells and induces Fas-dependent apoptosis. Mutations on the gene have been found in patients with severe developmental delay and epilepsy (Giorgio et al., 2015; Smigiel et al., 2016).

Lastly, CNVs in several other genes have been related to dyslexia and some of them are expected to be involved in the oscillopathic profile of this condition. *S100B* encodes a calcium-binding protein predominantly expressed in astrocytes, which is involved in neurite extension, stimulation of Ca^2+^ fluxes, and axonal proliferation, and ultimately, in synaptic plasticity and learning. Although *S100b* knockout mice show normal oscillation patterns in the neocortex and the hippocampus, they display a reduced gamma band (30-80 Hz) response in the hippocampus after seizure induction with kainic acid (Sakatani et al., 2008). Also, they kindle more rapidly and exhibit more severe seizures (Dyck et al., 2002). This abnormal response suggests that the S100B-related pathways may contribute to modulate brain oscillations and neural activities in specific conditions (Figure 1). Altered expression of *S100B* has been found in patients with mesial temporal epilepsy (Lu et al., 2010). *GABARAP* encodes a GABAA receptor-associated protein that clusters neurotransmitter receptors by promoting their interaction with the cytoskeleton. Specifically, GABARAP mediates inhibitory neural transmission via its interaction with KIF5A, which affects the neuronal surface expression of GABAA receptors: the conditional knockout of *Gabarap* in mice results in abnormal paroxysmal sharp waves in the hippocampus (Nakajima et al., 2012). Likewise, deletions of *GABARAP*, as part of a 2.3-Mb microdeletion of 17p13.2p13.1, have been found in patients with moderate mental retardation and intractable epilepsy (Komoike et al., 2010). *CNTNAP2* encodes a protein associated with K^+^ voltage-gated channels that regulates dendritic arborization and spine development (Anderson et al., 2012), axonogenesis in conjunction with ROBO factors (Banerjee et al., 2010), synaptogenesis (Dean et al., 2003), and brain connectivity and cerebral morphology (Scott-Van Zeeland et al., 2010; Tan et al., 2010; Dennis et al., 2011). Heterozygous mutations or CNVs of the gene have been related to different conditions entailing speech and language problems, including dyslexia (Peter et al., 2011), but also SLI (Newbury et al., 2011), child apraxia of speech (Worthey et al., 2013), variants of language delay and language impairment (Petrin et al., 2010; Sehested et al., 2010), ASD (Alarcón et al., 2008; Bakkaloglu et al., 2008), and language impairment in SZ (Poot, 2015). Non-pathogenic polymorphisms of CNTNAP2 seem to affect language development in the typically-developing population (Whalley et al., 2011; Whitehouse et al., 2011; Kos et al., 2012). Homozygous mutations or compound heterozygous CNVs of the gene result in epilepsy, and language and speech regression (Strauss et al., 2006; Marchese et al., 2016; Smogavec et al., 2016). Interestingly, rats and mice with homozygous deletions of *Cntnap2* show reduced spectral power in the alpha (9-12 Hz) range during wake (Thomas et al., 2017). *KANSL1* encodes a component of the NSL1 complex, which is involved in the acetylation of the nucleosomal histone H4, important for chromatin organization and gene transcription regulation. The gene is a candidate for Koolen-de Vrries syndrome (OMIM#610443), characterized by epilepsy, developmental delay, and moderate intellectual disability which impacts mostly on expressive language development (Koolen et al., 2016). Finally, *NSF* encodes a protein needed for vesicle-mediated transport in the Golgi apparatus and involved in synaptic function. One of the proteins that contribute to the formation of the soluble NSF attachment protein receptor complex (and ultimately, to the regulation of synaptic vesicle exocytosis) is SNAP25, which has been related to SZ and whose reduced levels are associated in mice with the occurrence of frequent spikes and diffuse network hyperexcitability, epileptiform discharges, and cognitive deficits (Corradini et al., 2014).

**Figure 1.**
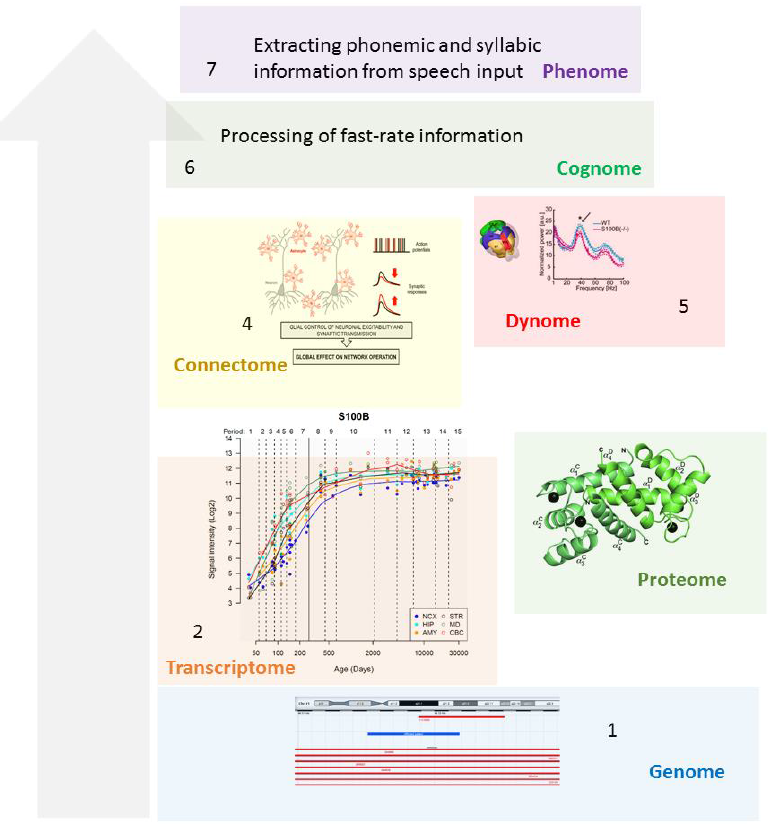
Observed and expected functional links between pathogenic changes in genes involved in brain rhythmicity and dyslexia-related features. These links are exemplified by S100B. As noted in the text, CNVs of this gene have been found in patients with the disease (1). The gene is expressed in several brain areas (2) and encodes a calcium-binding protein (3), which plays a role in neurite extension and axonal proliferation, and ultimately, in synaptic plasticity and learning (4). This last role might result in part from its effect on gamma oscillations in the hippocampus (5). As discussed in the text, altered levels of S100b affect gamma oscillations in this area. Because of the role of gamma rhythms in the processing of phonemic and syllabic information (6), mutations or dosage changes affecting S100B might account for some of the reading and spelling problems exhibited by dyslexics (7). The image of CNVs of the gene in (1) is from DECIPHER (https://decipher.sanger.ac.uk/). The expression data of S100B in the brain in (2) is from the Human Brain Transcriptome Database (http://hbatlas.org/). Six different brain regions are considered: the cerebellar cortex (CBC), the mediodorsal nucleus of the thalamus (MD), the striatum (STR), the amygdala (AMY), the hippocampus (HIP) and 11 areas of neocortex (NCX). The 3D view of the two S100B monomers in (3) is from Jensen et al. (2015) (Figure 1a, upper). The scheme in (4), showing the interaction between astrocytes and neurons, and the role of the former in synaptic response and neuronal network operation, is from Perea et al. (2014) (Figure 1). The reduced gamma peak (marked with an arrow) in the CA1 region of the hippocampus of S100b (-/-) mice after kainite injection in (5) is from Sakatani et al. 2008 (Figure 1D), whereas the schema of the mice hippocampus is from Kadakkuzha et al., 2015 (Figure 4A). Here, ‘genome’ refers to the whole set of genes related to dyslexia, ‘transcriptome’, to their RNA products, and ‘proteome’ to the protein molecules related to the disease. ‘Connectome’ refers to the wiring of the regions involved in sound and visual processing that are recruited for reading and spelling. ‘Dynome’ refers to the brain dynamics underlying (and permitting) these processes, in the line of Kopell et al., 2014 or Murphy, 2015. ‘Cognome’ refers to the basic cognitive operations underlying reading and spelling, in the line of Poeppel, 2012. Finally, ‘phenome’ refers to the discrete, language-specific activities involved in reading and spelling.

## 5. Conclusions

Next generation sequencing technologies have significantly increased the number of candidate genes for dyslexia. Molecular biology and neurobiology techniques are providing a better understanding of the functional roles played by these gene in the healthy and the dyslexic brains, both during development and in the adult state. Likewise, neuroimaging facilities are optimising our understanding of the dyslexic brain, in terms of its anatomical distinctive features and the way in which it processes sounds and visual cues during reading. Finally, we now know that cognitive functions usually result from complex interactions between close and distant brain areas, in which the coupling of brain oscillations at different frequency bands, plays a key role. Nonetheless, we still need to bridge the gap between the genetic variation found in dyslexics, the (abnormal) development of their brains, and the emergence of brain dysfunctions over time, which ultimately result in problems for reading and spelling. Ultimately, a better understanding of this complex etiopathogenesis of dyslexia is needed if we want to design better therapeutic approaches to the disease and improve the reading abilities of the affected people. In this paper we have tried to bridge this gap by constructing successful oscillatory endophenotypes of dyslexia and by trying to link them to the biological activity of genes that have been related to this condition. Although the number of these genes is still small, they map on cell functions (e.g. cation transportation), brain areas (e.g. temporal and auditory cortices), physiological aspects of brain function (e.g. GABA homeostasis), developmental processes (neuronal migration, axon guidance, neurite extension), and cognitive abilities (plasticity and learning) that are known to be impaired in dyslexics or in animal models of the disease, or that are hypothesised to play an important role in speech processing and reading tasks in the healthy population. Last but not least, we expect that our results contribute as well to the ongoing research program aimed at translating linguistic computation into a specific code of brain activity patterns).

